# Astrocytic RIPK3 exerts protective anti-inflammatory activity during viral encephalitis via induction of serpin protease inhibitors

**DOI:** 10.1101/2024.05.21.595181

**Authors:** Marissa Lindman, Irving Estevez, Eduard Marmut, Evan M. DaPrano, Tsui-Wen Chou, Kimberly Newman, Colm Atkins, Natasha M. O’Brown, Brian P. Daniels

## Abstract

Flaviviruses pose a significant threat to public health due to their ability to infect the central nervous system (CNS) and cause severe neurologic disease. Astrocytes play a crucial role in the pathogenesis of flavivirus encephalitis through their maintenance of blood-brain barrier (BBB) integrity and their modulation of immune cell recruitment and activation within the CNS. We have previously shown that receptor interacting protein kinase-3 (RIPK3) is a central coordinator of neuroinflammation during CNS viral infection, a function that occurs independently of its canonical function in inducing necroptotic cell death. To date, however, roles for necroptosis-independent RIPK3 signaling in astrocytes are poorly understood. Here, we use mouse genetic tools to induce astrocyte-specific deletion, overexpression, and chemogenetic activation of RIPK3 to demonstrate an unexpected anti-inflammatory function for astrocytic RIPK3. RIPK3 activation in astrocytes was required for host survival in multiple models of flavivirus encephalitis, where it restricted neuropathogenesis by limiting immune cell recruitment to the CNS. Transcriptomic analysis revealed that, despite inducing a traditional pro-inflammatory transcriptional program, astrocytic RIPK3 paradoxically promoted neuroprotection through the upregulation of serpins, endogenous protease inhibitors with broad immunomodulatory activity. Notably, intracerebroventricular administration of SerpinA3N in infected mice preserved BBB integrity, reduced leukocyte infiltration, and improved survival outcomes in mice lacking astrocytic RIPK3. These findings highlight a previously unappreciated role for astrocytic RIPK3 in suppressing pathologic neuroinflammation and suggests new therapeutic targets for the treatment of flavivirus encephalitis.

## Introduction

Flaviviruses pose an escalating threat to public health, driven by the expanding habitats of their arthropod vectors (*1–3*). Clinically important flaviviruses include West Nile virus (WNV), Zika virus (ZIKV), Tick-borne encephalitis virus (TBEV), and Japanese encephalitis virus (JEV), all of which can invade and infect the central nervous system (CNS) (*4–7*). Neuroinvasive flavivirus infections are often fatal, and survivors frequently face persistent neurological sequalae long after the resolution of infection (*8–10*). Thus, identifying mechanisms of neuropathogenesis and host protection during flavivirus encephalitis is essential for the development of targeted therapeutics to treat and manage neuroinvasive infection (*11, 12*).

Following the invasion of flaviviruses into CNS tissues, neural cells mount robust immune responses that are essential for controlling viral spread (*13–15*). These responses are critical for cell-intrinsic restriction of viral replication, as well as for recruiting peripheral immune cells that coordinate the eradication of infection (*16, 17*). However, while peripheral immune cell recruitment to the CNS is necessary for viral clearance, infiltrating leukocytes are also capable of driving significant immunopathology and bystander injury of uninfected neural cells (*18–20*). Thus, the complex neuroimmune response to neuroinvasive viral infection must be tightly regulated to promote neuroprotection while limiting immunopathogenesis.

Recent work from our laboratory and others has identified a significant role for receptor interacting protein kinase-3 (RIPK3) in coordinating neuroimmune responses to flavivirus infections in the CNS (*21, 22*). RIPK3 is canonically associated with a form of lytic programmed cell death known as “necroptosis,” although a growing body of work has now defined extensive functions for RIPK3 that occur independently of cell death (*23–28*). During flavivirus encephalitis, we have shown that RIPK3 signaling in neurons drives a transcriptional program that includes a broad variety of pro-inflammatory and antiviral effector genes that work to control infection without inducing necroptosis (*29–31*). However, roles for RIPK3 in non-neuronal cells during neuroinvasive viral infections remain largely unexplored. Astrocytes are a highly abundant glial cell type that serve complex roles in both CNS homeostasis and disease. As integral components of the neurovascular unit, they regulate the integrity of the blood-brain barrier (BBB) and exert regulatory control over the recruitment and infiltration of leukocytes into the CNS parenchyma during neuroinflammatory disease states (*32*). The crucial anatomical positioning of these cells at the BBB and their central function in the regulation of neuroinflammation suggests the possibility of unique, cell type-specific functions for RIPK3 in astrocytes during flavivirus infections.

In this study, we demonstrate unexpected protective functions for astrocytic RIPK3 signaling in restricting neuroinflammation. Contrary to its predominantly proinflammatory function in neurons (*29–31*), we show that RIPK3 activation in astrocytes suppresses neuropathogenesis by limiting immune cell recruitment to the CNS during flavivirus encephalitis. Using conditional *in vivo* expression tools, including an inducible chemogenetic RIPK3 activation system, we show that the RIPK3-dependent transcriptional program in astrocytes is enriched for serpins, a family of endogenous serine protease inhibitors with immunomodulatory activity. Notably, mice deficient in astrocytic *Ripk3* exhibited significant BBB disruption and enhanced leukocyte infiltration into the CNS. However, reconstitution of SerpinA3N in the CNS of acutely infected mice preserved BBB integrity, reduced CNS leukocyte infiltration, and prevented fatal outcomes during flavivirus encephalitis. These findings underscore previously unappreciated, cell-type specific functions for RIPK3 in astrocytes in promoting host protection during CNS viral infection.

## Results

### Astrocytic RIPK3 restricts flavivirus pathogenesis but not cell-intrinsic viral replication

To determine whether RIPK3 signaling in astrocytes had a role in neuropathogenesis during CNS flavivirus infection, we generated mice with astrocyte-specific deficiency of *Ripk3* by crossing mice in which exons 2 and 3 of the endogenous *Ripk3* locus are flanked by loxP sites (*33*) (*Ripk3*^fl/fl^) to a line expressing tamoxifen-inducible Cre-recombinase under the control of the *Aldh1l1* promoter (*Aldh1l1* Cre/ERT2, shorthanded as *Aldh1l1* Cre throughout this manuscript). We subjected these animals to several models of flavivirus encephalitis, including intracranial infection with an ancestral African strain of ZIKV (ZIKV-MR766, Uganda 1947) or a contemporary Asian lineage strain (ZIKV-Fortaleza, Brazil 2015). We also used a subcutaneous infection model with a neurovirulent North American strain of WNV (WNV-WN02-Bird114, Texas 2002) (Figure 1A). Notably, mice lacking astrocytic RIPK3 expression (*Ripk3*^fl/fl^ *Aldh1l1* Cre^+^) exhibited accelerated and enhanced mortality compared to littermate controls in all three infection models (Figure 1B-D). Mice also exhibited more rapid onset of clinical disease in all three models, as evidenced by earlier and more pronounced weight loss in the first six days of infection (Figure 1B-D), suggesting that astrocytic RIPK3 restricts pathogenesis in these models. As a complimentary approach, we also used mice harboring astrocyte-specific overexpression of RIPK3. This line expresses a chimeric version of RIPK3 (RIPK3-2xFV) under the control of a lox-STOP-lox element in the *Rosa26* locus. RIPK3-2xFV features two FKBP^F36V^ domains that facilitate enforced oligomerization following treatment with a dimerization drug. This construct leaves the endogenous *Ripk3* locus intact, and thus this mouse line can be used as either a cell type-specific overexpression system or an enforced chemogenetic activation system in cell types of interest in vivo (*29, 30, 34*). Notably, mice with astrocyte-specific overexpression of RIPK3 (*Ripk3*-2xFV^fl/fl^ *Aldh1l1* Cre^+^) exhibited significant amelioration of disease following intracranial challenge with ZIKV-MR766 (Figure 1E), further supporting a protective function for astrocytic RIPK3 during flavivirus infection.

**Figure 1:**
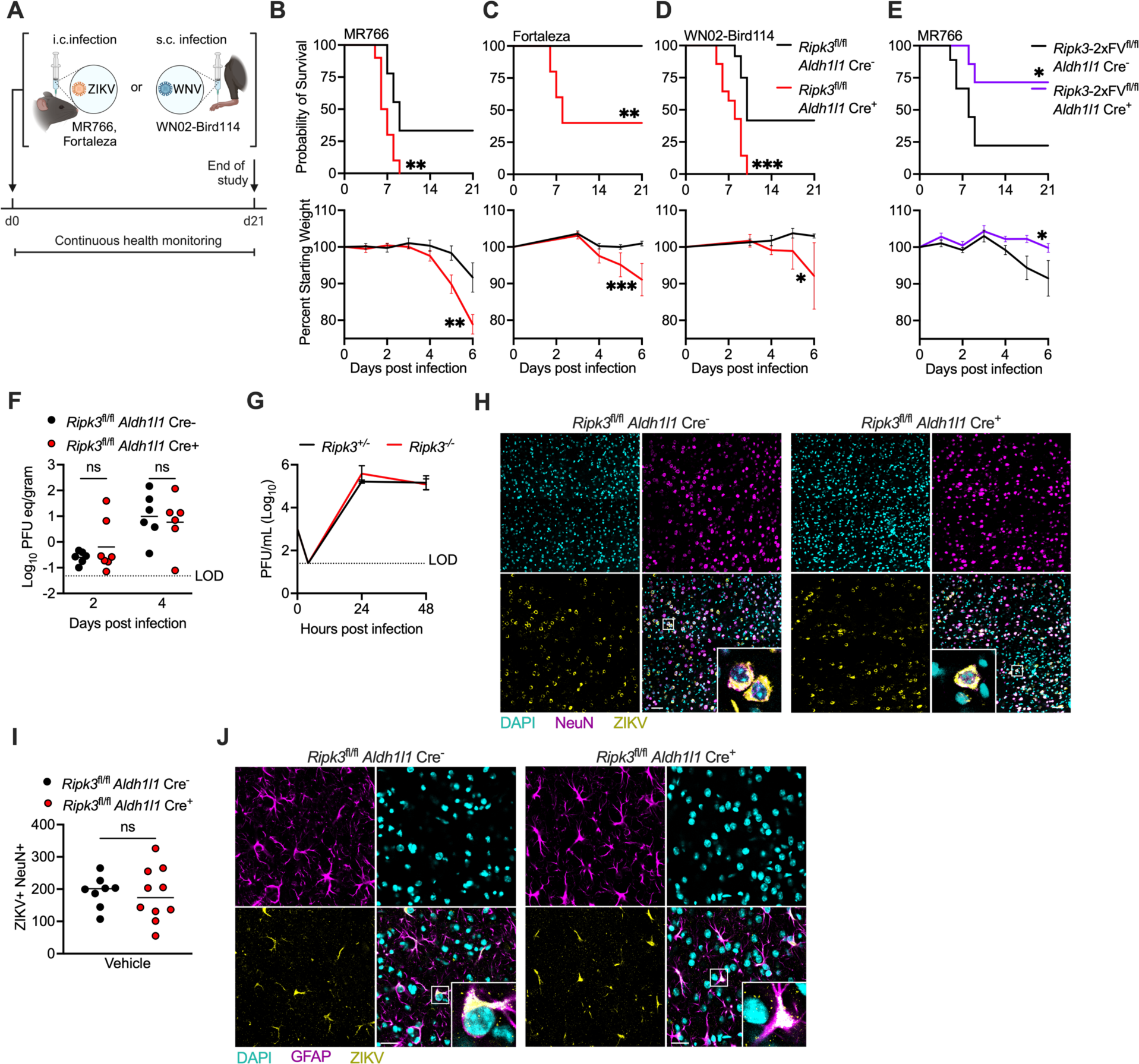
Astrocytic RIPK3 restricts flavivirus pathogenesis but not cell-intrinsic viral replication. (**A**) Schematic illustrating survival studies in which mice underwent either intracranial (i.c.) ZIKV or subcutaneous (s.c.) WNV infections. (**B-E**) Survival and weight measurements in mice of indicated genotypes following i.c. ZIKV-MR766 infection (B), i.c. ZIKV-Fortaleza infection (C) infection, s.c. WNV-WN02-Bird114 infection (D), or i.c. ZIKV-MR766 infection (E). n=5-14 mice/group. (**F**) qRT-PCR analysis of ZIKV-MR766 genome copies in whole brain homogenates derived from mice of indicated genotypes, two or four days following i.c. ZIKV-MR766 infection. Data are expressed as plaque forming unit equivalents (PFU eq)/gram of brain tissue. (**G**) Multistep growth curve analysis of ZIKV-MR766 replication in primary cortical astrocytes derived from mice of indicated genotypes. Viral titers were assessed via plaque assay. N=4 replicates/group. (**H**) Immunohistochemical (IHC) staining of NeuN (magenta), DAPI (cyan), and pan-flavivirus E protein (yellow) staining in cortical brain tissue derived from mice of indicated genotypes five days following i.c. ZIKV-MR766 infection. Images are 2×2 tiled composites at 20x magnification. Scale bar = 20μm. (**I**) Quantification of cells exhibiting colocalization of NeuN and pan-flavivirus E protein in cortical brain tissue derived from mice described in (H). (J) IHC staining of GFAP (magenta), DAPI (cyan), and pan-flavivirus E protein (yellow) in mice as described in (H). Scale bar = 20μm. Images are at 40x magnification. *p<0.05, **p < 0.01, ***p < 0.001. Error bars represent SEM.

We next questioned whether astrocytic RIPK3 promotes host survival through direct suppression of flavivirus replication and brain viral burden. Analysis of ZIKV RNA in brain homogenates showed no differences between *Ripk3*^fl/fl^ *Aldh1l1* Cre^+^ mice and littermate controls following ZIKV-MR766 infection (Figure 1F). Multistep growth curve analysis also showed no difference in ZIKV-MR766 replication in primary astrocytes derived from *Ripk3*^−/−^ mice compared to cultures derived from heterozygous littermate controls (Figure 1G). We also saw no differences in replication of either ZIKV-MR766 or WNV-WN02-Bird114 in primary astrocytes treated with GSK872, an inhibitor of RIPK3 kinase function (Figure S1A-B). We also observed no impact of *Mlkl* deficiency on ZIKV-MR766 replication (Figure S1C), or of either *Ripk3* or *Mlkl* deficiency on astrocyte viability following infection (Figure S1D), confirming that RIPK3 does not control ZIKV infection via induction of necroptosis, consistent with our previous work (*29*). Immunohistochemical (IHC) analysis of infected cells in the cerebral cortex of animals infected with ZIKV-MR766 also showed no impact of astrocytic RIPK3 deletion on numbers of infected neurons (Figure 1H-I), which represent the vast majority of infected cells during flavivirus encephalitis (*18, 35, 36*). While infection of astrocytes is more sporadic in this model and therefore difficult to systematically quantify, we also did not observe qualitative differences in the location or numbers of infected astrocytes between genotypes (Figure 1J). Together, these data strongly suggest that the protective benefit of astrocytic RIPK3 was not achieved through direct modulation of infection or viral replication in the CNS.

### Activation of astrocytic RIPK3 drives proinflammatory gene expression

Given our observation of a protective function for endogenous RIPK3 activity during flavivirus encephalitis, we next questioned whether exogenous activation of RIPK3 would have therapeutic efficacy. We thus infected mice expressing the chemogenetically activatable RIPK3-2xFV protein in astrocytes (*Ripk3*-2xFV^fl/fl^ *Aldh1l1* Cre^+^) with ZIKV-MR766. Mice were treated on day three post infection with B/B homodimerizer (B/B), which drives dimerization and activation of RIPK3-2xFV, or a vehicle control solution (Figure 2A). Chemogenetic activation of RIPK3 conferred significant protection from intracranial ZIKV-MR766 challenge, as indicated by essentially complete amelioration of mortality and weight loss in the B/B-treated group (Figure 2B). To better understand mechanisms of neuroprotection induced by astrocytic RIPK3, we performed magnetic activated cell sorting (MACS) of ACSA2^+^ astrocytes 24 hours following B/B administration in *Ripk3*-2xFV^fl/fl^ *Aldh1l1* Cre^+^ mice or their Cre^−^ littermate controls (Figure 2C). Sorted astrocytes were subjected to bulk RNA-sequencing (RNAseq). Astrocytes derived from *Ripk3*-2xFV^fl/fl^ *Aldh1l1* Cre^+^ mice exhibited robust transcriptomic changes compared to those derived from controls, as evidenced by distinct clustering in principal component analysis (PCA) (Figure 2D). Chemogenetic RIPK3 activation in astrocytes resulted in 3,662 significantly differentially expressed genes (DEGs) in total, including 1,691 downregulated DEGs and 1,971 upregulated DEGs (Figure 2E). Gene ontology (GO) enrichment analysis revealed significant overrepresentation of terms related to immune cell trafficking and inflammatory signaling, consistent with previous work from our group and others establishing RIPK3 as a key coordinator of neuroinflammation (Table S1). Further analysis of selected enriched GO terms revealed predominantly upregulated expression of genes associated with Type I interferon signaling (Figure 2F), cytokine responses (Figure 2G), and T cell mediated immunity (Figure 2H), consistent with an overall proinflammatory transcriptomic signature in the setting of RIPK3 activation. We also confirmed that endogenous *Ripk3* was required for induction of inflammatory chemokines and interferon stimulated genes (ISGs) in primary astrocytes infected with ZIKV-MR766 (Figure 2I). Together, these findings demonstrate that both chemogenetic and infection-induced activation of RIPK3 drives proinflammatory gene expression in astrocytes.

**Figure 2:**
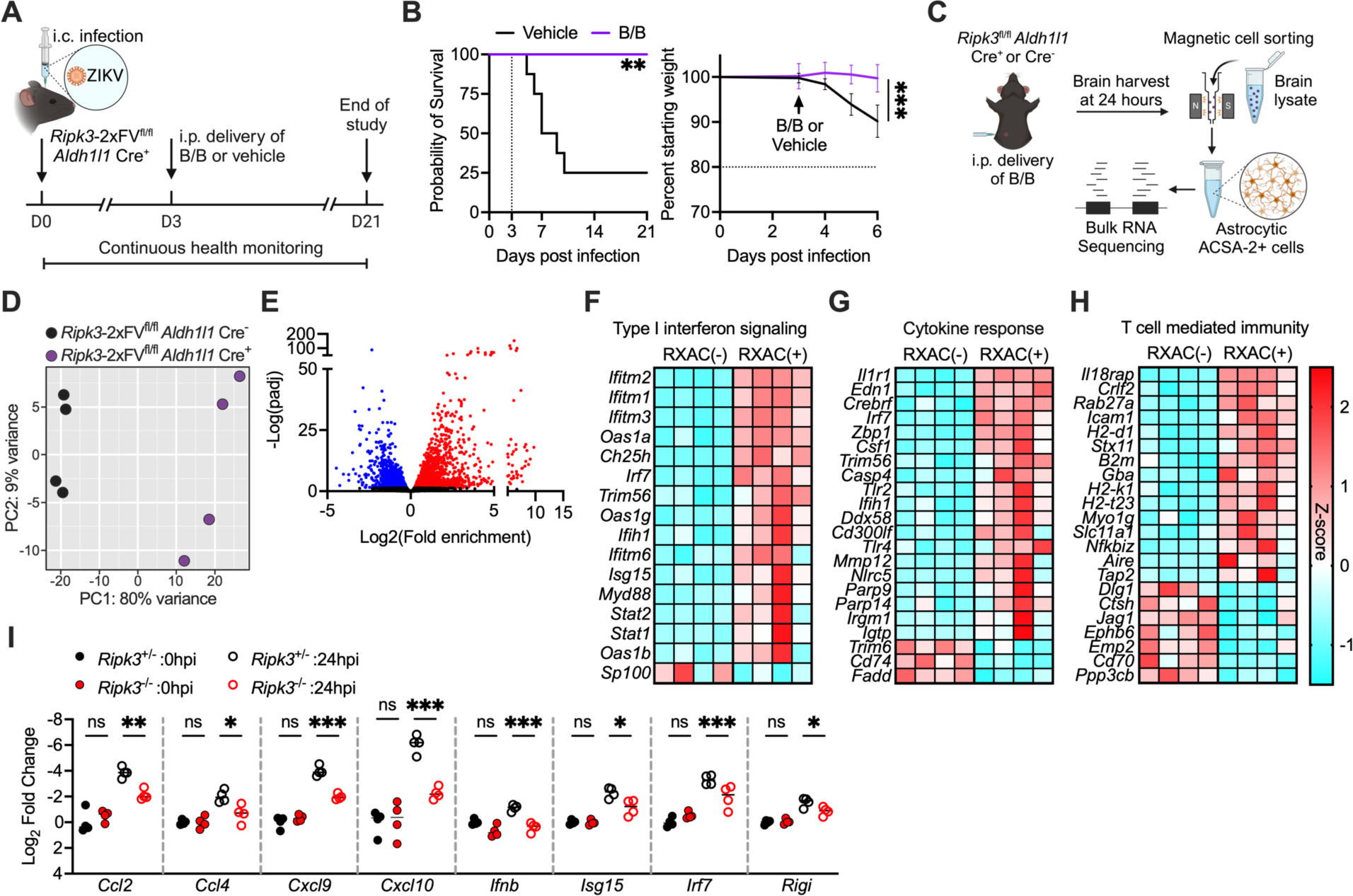
Activation of astrocytic RIPK3 drives proinflammatory gene expression. (**A**) Schematic illustrating survival studies in which mice expressing a chemogenetically activatable form of RIPK3 (*Ripk3*-2xFV) received intraperitoneal (i.p.) administration of homodimerizer ligand (B/B) three days following high dose (10^3^ PFU) intracranial (i.c.) ZIKV-MR766 infection. (**B**) Survival and weight measurements in *Ripk3*-2xFV^fl/fl^ *Aldh1l1* Cre^+^ mice treated as described in (A). (**C**) Schematic illustrating magnetic activated cell sorting (MACS) of ACSA-2^+^ astrocytes derived from mice of indicated genotypes 24 hours following i.p. administration of B/B. (**D-E**) Principal component analysis (PCA) (D) and volcano plot depicting significant differentially expressed genes (E) in bulk RNA sequencing data derived from ACSA-2^+^ astrocytes as described in (C). Transcripts with significant differential expression (>1.5-fold change, adj. P < 0.05) are highlighted. Downregulated transcripts are shown in blue and upregulated transcripts are shown in red. (**F-H**) Heatmap depicting significant differentially expressed genes within indicated gene ontology terms. (I) qRT-PCR analysis of indicated inflammatory chemokines and interferon stimulated genes in primary astrocytes derived from mice of indicated genotypes 0 or 24 hours following ZIKV-MR766 infection. *p<0.05, **p < 0.01, ***p < 0.001. Error bars represent SEM.

### Astrocytic RIPK3 suppresses CNS leukocyte infiltration during flavivirus encephalitis

We previously demonstrated a role for RIPK3 signaling in neurons to drive recruitment of antiviral leukocytes into the CNS through the induction of inflammatory chemokine expression (*30*). Given that activation of RIPK3 in astrocytes also appeared to drive chemokine expression, we questioned if astrocytic RIPK3 similarly supported leukocyte recruitment to the CNS. We thus performed flow cytometric analysis of CNS leukocytes on day 4 following intracranial ZIKV-MR766 infection (Figure 3A). Surprisingly, we observed enhanced frequencies of CD45^hi^ infiltrating cells in *Ripk3*^fl/fl^ *Aldh1l1* Cre^+^ mice compared to littermate controls following infection (Figure 3B). In contrast, frequencies of CD45^int^ CD11b^+^ resident microglia were unchanged between genotypes (Figure 3C). Further profiling revealed enhanced infiltration of several myeloid cell populations in *Ripk3*^fl/fl^ *Aldh1l1* Cre^+^ mice, including CD11b^+^ Ly6c^+^ monocytes and CD11b^+^ F4/80^+^ macrophages, as well as CD11c^+^ MHCII^+^ dendritic cells (Figure 3D). In the lymphocyte compartment, we also observed significant increases in both CD8^+^ and CD4^+^ T cell subsets, as well as NK1.1^+^ natural killer cells (Figure 3E-F). Importantly, these changes were not due to baseline differences in CNS leukocyte populations between genotypes, as we did not observe differences in any CNS leukocyte subset in mock-infected animals (Figure S2A, Figure 3C, D, and F). We also did not observe differences in non-CNS immune responses, including leukocyte frequencies in the spleen (Figure S2B-C). These data suggest that, despite promoting a broad set of traditionally pro-inflammatory genes, RIPK3 activity in astrocytes nevertheless exerts an *anti-inflammatory* effect on CNS leukocyte infiltration during flavivirus encephalitis.

**Figure 3:**
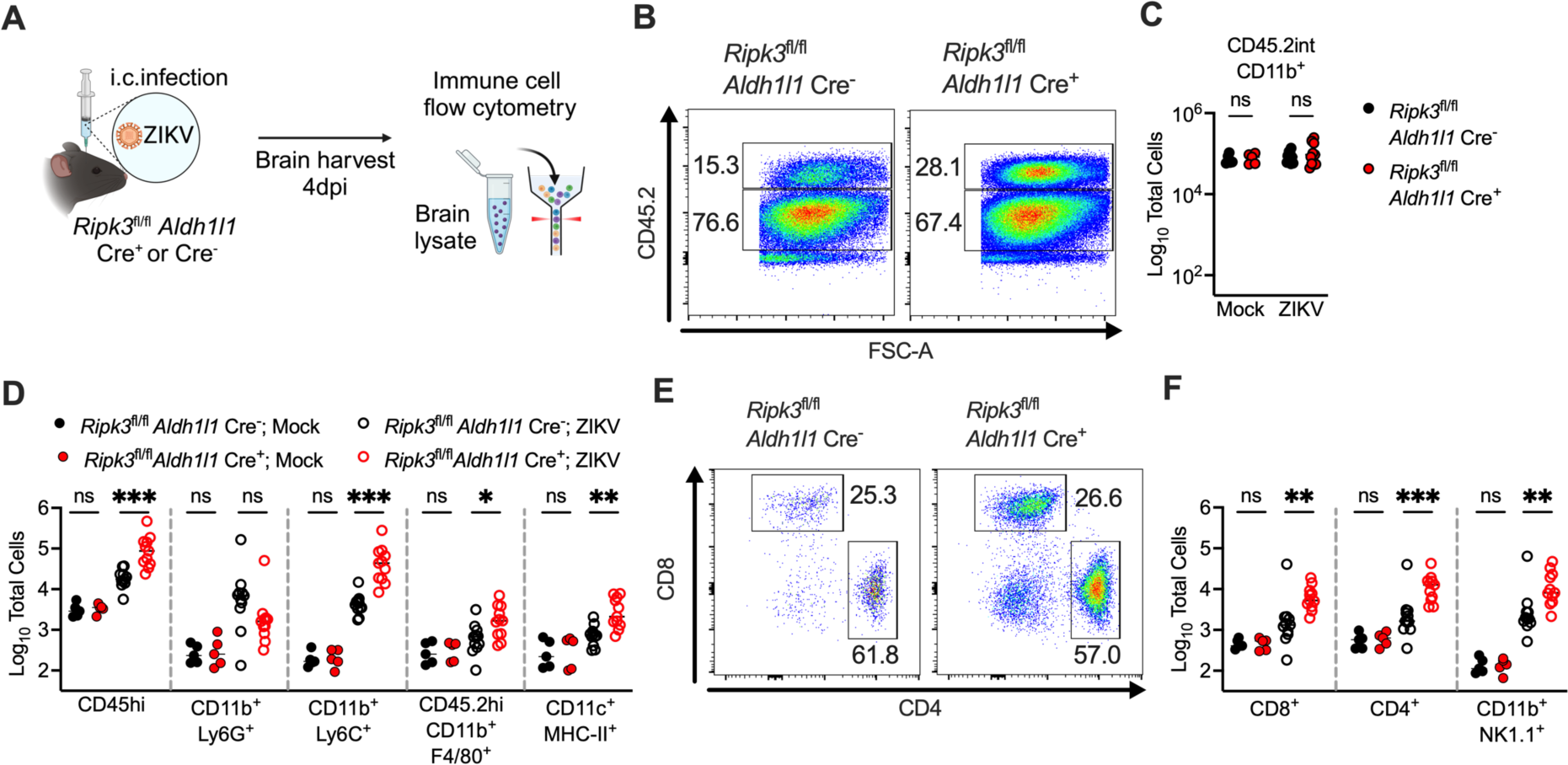
Astrocytic RIPK3 suppresses CNS leukocyte infiltration during flavivirus encephalitis. (**A**) Schematic illustrating flow cytometric analysis of CNS leukocytes derived from mice of indicated genotypes four days following intracranial (i.c.) ZIKV-MR766 infection. (**B**) Representative flow cytometry plots depicting resident (CD45.2^int^) or infiltrating (CD45.2^hi^) CNS leukocytes derived from mice infected with ZIKV-MR766 as described in (A). (**C**) Total numbers of CD45^int^ CD11b^+^ resident microglia in brain tissue derived from mice of indicated genotypes following ZIKV-MR766 infection. (**D**) Total numbers of indicated myeloid cell populations in brain tissue derived from mice of indicated genotypes following ZIKV-MR766 infection. (**E**) Representative flow cytometry plots depicting CD4^+^ T cells and CD8^+^ T cells derived as described in (A). (**F**) Total numbers of CD8^+^ T cells, CD4^+^ T cells, and NK1.1^+^ natural killer cells in brain tissue derived from mice of indicated genotypes following ZIKV-MR766 infection. *p<0.05, **p < 0.01, ***p < 0.001.

### Astrocytic RIPK3 promotes a complex immunologic transcriptional program, including multiple Serpin protease inhibitors

Given these paradoxical findings, we returned to our transcriptomic analysis to assess how astrocytic RIPK3 influences gene pathways associated with anti-inflammatory signaling and immunoregulation. Notably, we observed enrichment of several relevant gene ontology terms in our dataset, including “Inhibition of inflammatory response,” “Inhibition of lymphocyte activation,” and others (Figure 4A). Analysis of expression patterns within these enriched GO terms revealed complex patterns of both up- and down-regulated expression, suggesting that RIPK3 likely exerts more nuanced control over inflammatory signaling in astrocytes than has been appreciated previously (Figure 4B, FigureS3). Among the DEGs induced by RIPK3 activation in astrocytes, we noticed that serpins were particularly enriched, representing 5 of the top 25 overall upregulated DEGs (Figure 4C). Serpins are a family of endogenous serine protease inhibitors that exert diverse immunoregulatory functions, including inhibition of leukocyte proteases and matrix-metalloproteinases (MMPs) that support inflammation (*37, 38*). These molecules thus stood out to us as a promising candidate mechanism of immunoregulation downstream of astrocytic RIPK3. In total, chemogenetic activation of RIPK3 significantly altered the expression of 15 serpin molecules (Figure 4D). We also confirmed that endogenous *Ripk3* was required for induction of serpin expression in primary mouse astrocytes infected with ZIKV-MR766 (Figure 4E). We have previously shown that RIPK3 activation in astrocytes regulates transcription through induction of the inflammatory transcription factor NFκB (*39, 40*). As many serpins contain κB elements in their promoters (*41–43*), we tested a requirement for NFκB in astrocytic serpin expression using inhibitors of NFκB signaling, including BAY 11-7085 (an inhibitor of IKK activity) and JSH-23 (an inhibitor of NFκB p65 nuclear translocation). Both inhibitors abrogated induction of serpin expression following ZIKV-MR766 infection (Figure 4E). Together, these data identify serpins as candidate immunoregulatory molecules induced downstream of RIPK3 and NFκB in astrocytes.

**Figure 4:**
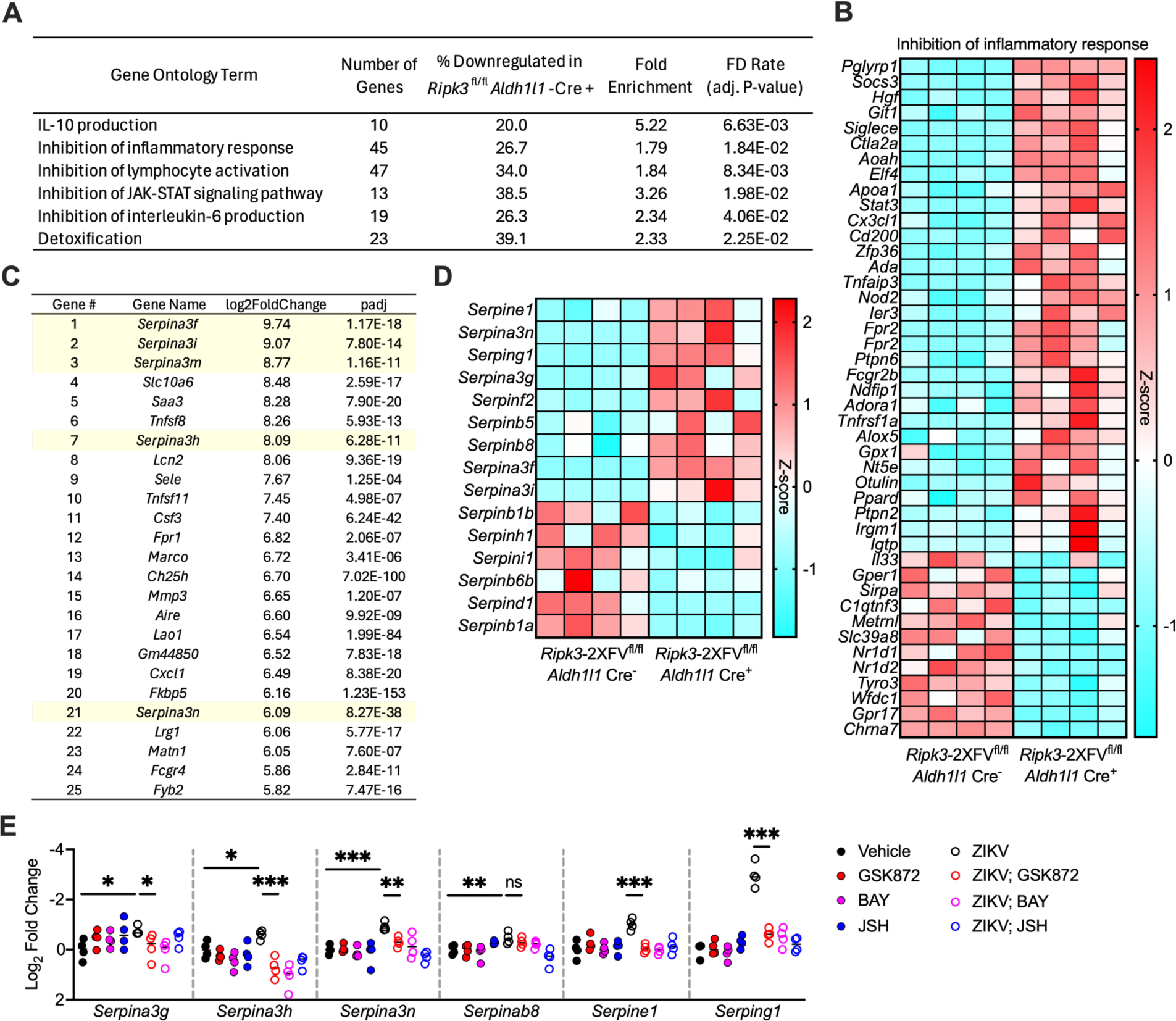
Astrocytic RIPK3 promotes a complex immunologic transcriptional program, including multiple Serpin protease inhibitors. (**A**) Gene ontology enrichment analysis of terms related to anti-inflammatory signaling and immunoregulation, derived from RNAseq analysis comparing ACSA-2^+^ astrocytes sorted from *Ripk3*-2xFV^fl/fl^ *Aldh1l1* Cre^+^ mice compared to astrocytes sorted from Cre^−^ littermate controls 24 hours following B/B treatment. (**B**) Heatmap depicting significant differentially expressed genes within indicated gene ontology term. (**C**) Top 25 differentially expressed genes induced by chemogenetic RIPK3 activation in ACSA-2^+^ astrocytes. (**D**) Heatmap depicting all significantly differentially expressed serpin genes induced by chemogenetic RIPK3 activation in astrocytes. (**E**) qRT-PCR analysis of indicated serpin genes in primary astrocytes derived from WT mice. Astrocytes were pretreated for two hours with indicated inhibitors then infected with ZIKV-MR766. Gene expression was measured 24 hours post infection. *p<0.05, **p < 0.01, ***p < 0.001.

### SerpinA3N is neuroprotective in models of flavivirus encephalitis

Among the differentially expressed serpin molecules in our transcriptomic analysis, we noted SerpinA3N due to numerous recent reports of its high expression in astrocytes and involvement in a wide variety of neurologic diseases (*41*). We confirmed that SerpinA3N protein expression is elevated in whole brain homogenates in control animals infected with ZIKV-MR766, while this increase was reduced in *Ripk3*^fl/fl^ *Aldh1l1* Cre^+^ mice (Figure 5A). We next questioned whether reconstitution of SerpinA3N protein in the CNS of *Ripk3*^fl/fl^ *Aldh1l1* Cre^+^ mice would impact neuropathogenesis. We thus performed intracerebroventricular (ICV) administration of recombinant murine SerpinA3N in mice two days following intracranial ZIKV-MR766 infection (Figure 5B). As SerpinA3N is a potent inhibitor of serine protease activity, we first measured overall levels of proteolysis in the brain using an assay that measures endogenous collagenase/gelatinase activity via cleavage of a fluorogenic gelatin substrate. Brain homogenates from *Ripk3*^fl/fl^ *Aldh1l1* Cre^+^ mice exhibited significantly enhanced collagenase/gelatinase activity compared to littermate controls; however, this effect was rescued following ICV injection of SerpinA3N (Figure 5C). Among the serine proteases inhibited by SerpinA3N in the brain are matrix metalloproteinases and other enzymes that degrade extracellular matrix and tight junction (TJ) molecules that support BBB function. We thus performed immunostaining for the BBB-specific TJ molecule Claudin-5 and its adapter ZO-1, which neatly colocalize in brain vascular endothelia under homeostatic conditions. We observed significant perturbation of Claudin-5 and ZO-1 colocalization in *Ripk3*^fl/fl^ *Aldh1l1* Cre^+^ mice compared to controls, driven primarily by loss of ZO-1 expression (Figure 5D-E). However, this effect could also be prevented via administration of exogenous SerpinA3N. Given these confirmations that exogenous SerpinA3N treatment could rescue changes in brain protease activity and TJ formation in animals lacking astrocytic RIPK3, we next questioned whether this intervention would have clinical efficacy in models of flavivirus encephalitis. We thus performed ICV SerpinA3N administration at time points preceding the onset of clinical disease in the setting of intracranial ZIKV-MR766 infection or subcutaneous WNV-WN02-Bird114 infection (Figure 5F). Injection of SerpinA3N at day two post intracranial ZIKV-MR766 infection significantly ameliorated both weight loss and overall animal mortality in *Ripk3*^fl/fl^ *Aldh1l1* Cre^+^ mice (Figure 5G). We observed similar results using two injections of SerpinA3N at days five and seven following subcutaneous WNV-WN02-Bird114 infection (Figure 5H). We note that two injections were used in this model due to the delayed and more protracted course of disease that occurs following subcutaneous (rather than intracranial) infection. Together, these data confirm that SerpinA3N confers neuroprotection during flavivirus encephalitis and suggest that diminished expression of SerpinA3N is a mechanism of enhanced disease burden in mice lacking astrocytic RIPK3.

**Figure 5:**
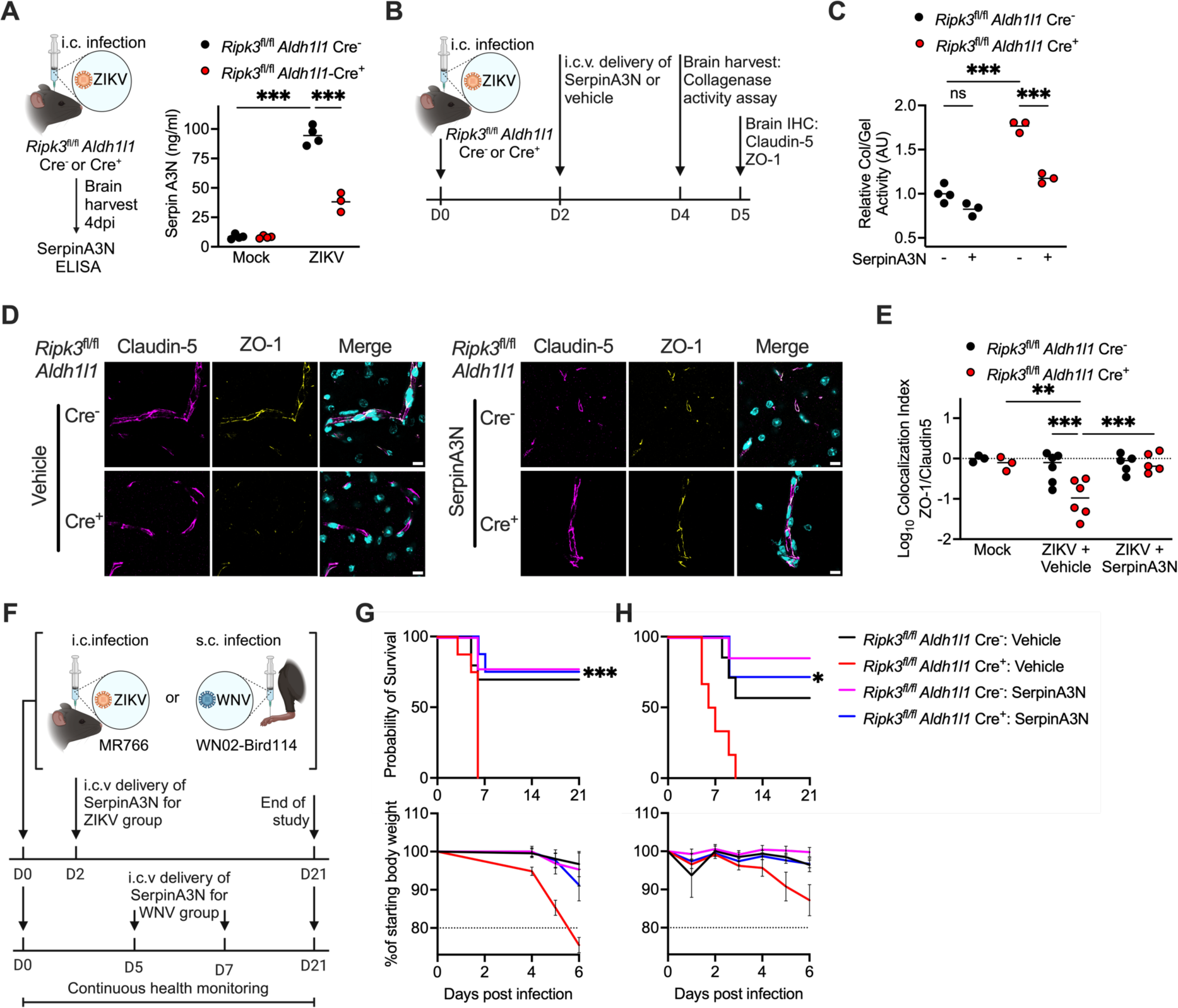
SerpinA3N is neuroprotective in models of flavivirus encephalitis. (**A**) SerpinA3N concentrations (measured via ELISA) in whole brain homogenates derived from mice of indicated genotypes four days following intracranial (i.c.) ZIKV-MR766 infection. (**B**) Schematic illustrating timeline of i.c. ZIKV-MR766 infection and i.c.v. treatment with recombinant SerpinA3N, prior to various endpoint readouts. (**C**) Collagenase/gelatinase activity assay in whole brain homogenates derived from mice of indicated genotypes following ZIKV-MR766 infection and SerpinA3N treatment, as described in (B). (**D**) Immunohistochemical (IHC) staining of Claudin5 (magenta), Zo-1 (yellow), and DAPI (cyan) in cortical brain tissue of mice of indicated genotypes following ZIKV-MR766 infection and SerpinA3N treatment, as described in (B). Scale bar = 10μm. Images are at 63x magnification. (**E**) Colocalization index of Claudin5 and Zo-1 staining in cortical brain sections described in (D). (**F**) Schematic illustrating survival studies in which mice received i.c. ZIKV-MR766 infection or subcutaneous (s.c.) WNV-WN02-Bird114 infection, followed by i.c.v. administration of recombinant SerpinA3N at indicated intervals following infection. (**G-H**) Survival and weight measurements in mice of indicated genotypes following ZIKV-MR766 infection (G) or WNV-WN02-Bird114 infection (H) with or without i.c.v. SerpinA3N treatment, as described in (F). n=6-10 mice/group. *p<0.05, **p < 0.01, ***p < 0.001. Error bars represent SEM.

### SerpinA3N suppresses deleterious neuroinflammation during flavivirus encephalitis

We next assessed potential mechanisms by which SerpinA3N ameliorates pathogenesis during flavivirus encephalitis. Notably, recombinant SerpinA3N treatment did not have a direct antiviral effect against either ZIKV-MR766 or WNV-WN02-Bird114 in astrocytes *in vitro*, suggesting that, like RIPK3, SerpinA3N does not limit pathogenesis through restriction of viral replication (Figure S4A-B). In addition to inhibiting MMPs, SerpinA3N is known to inhibit the activity of several proteases involved in inflammatory signaling, including cathepsins, leukocyte elastase, and granzyme B (*44*). We thus questioned whether lack of SerpinA3N might be a mechanism of enhanced neuroinflammation in *Ripk3*^fl/fl^ *Aldh1l1* Cre^+^ mice during flavivirus infection. Remarkably, ICV delivery of exogenous SerpinA3N two days following intracranial ZIKV infection significantly ameliorated the enhanced CNS leukocyte infiltration observed in *Ripk3*^fl/fl^ *Aldh1l1* Cre^+^ mice (Figure 6A). This effect occurred across several leukocyte subsets, including both CD4^+^ and CD8^+^ T cells, NK cells, and monocytes (Figure 6B). Among these subsets, previous work has established pathogenic roles for infiltrating CD8^+^ T cells, in particular, in driving host mortality during flavivirus encephalitis (*18, 19*). We therefore questioned whether the enhanced infiltration of CD8^+^ T cells was a mechanism underlying the enhanced pathogenesis observed in mice lacking astrocytic RIPK3 following flavivirus infection. To do so, we depleted CD8^+^ T cells via administration of an anti-CD8 neutralizing antibody two days following intracranial ZIKV-MR766 infection (Figure 6C). We confirmed complete depletion of circulating CD8s two days later (day four post infection) (Figure S5A-B). CD8 depletion significantly ameliorated the enhanced mortality and weight loss observed in *Ripk3*^fl/fl^ *Aldh1l1* Cre^+^ mice following infection, confirming a pathogenic role for this cell type in the absence of astrocytic RIPK3 (Figure 6D). This diminishment of animal mortality in CD8-depleted *Ripk3*^fl/fl^ *Aldh1l1* Cre^+^ animals was also associated with decreased signs of neuroinflammation, as indicated by diminished immunoreactivity for GFAP in the cerebral cortex (Figure 6E-F). Together, these data support a model in which astrocytic RIPK3 suppresses immunopathology during flavivirus encephalitis through the induction of serpin protease inhibitors, such as SerpinA3N, which maintain BBB integrity and restrict deleterious recruitment of pathogenic CD8^+^ T cells to the brain.

**Figure 6:**
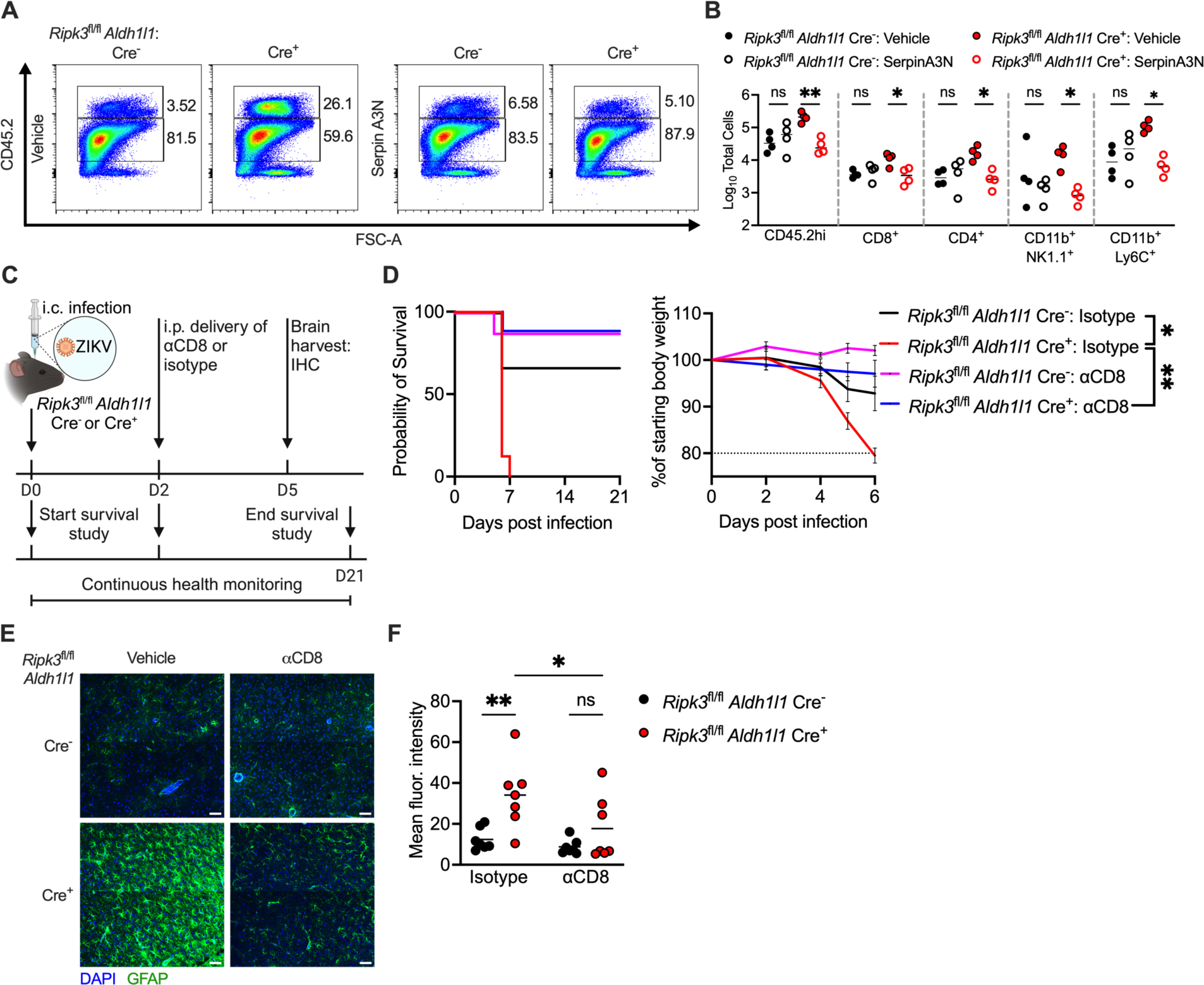
SerpinA3N suppresses deleterious neuroinflammation during flavivirus encephalitis. (**A**) Representative flow cytometry plots depicting resident (CD45.2^int^) or infiltrating (CD45.2^hi^) CNS leukocytes derived from mice of indicated genotypes infected with ZIKV-MR766 with or without i.c.v. treatment with SerpinA3N two days following infection. Flow cytometry was performed four days following infection. (**B**) Total numbers of indicated leukocyte populations in brain tissue derived from mice described in (A). (**C**) Schematic illustrating survival study and immunohistochemical (IHC) analysis in which mice received intraperitoneal administration of an anti-CD8 neutralizing antibody (αCD8) two days following intracranial (i.c.) ZIKV-MR766 infection. (**D**) Survival and weight measurements in mice of indicated genotypes following ZIKV-MR766 infection with or without depletion of CD8^+^ T cells, as described in (C). (**E**) IHC analysis of GFAP (green) and DAPI (blue) in cortical brain tissue derived from mice of indicated genotypes treated as described in (C). Images are 2×2 tiled composites at 20x magnification. Scale bar = 50μm. (**F**) Mean fluorescence intensity of GFAP staining in cortical brain tissue derived from mice of indicated genotypes and treated as described in (C). *p<0.05, **p < 0.01. Error bars represent SEM.

## Discussion

Recent work has broadened our understanding of RIPK3 signaling to include functions beyond its traditional association with necroptosis. To date, these cell death-independent functions have primarily been pro-inflammatory in nature. Within the CNS, we and others have shown that RIPK3 signaling is a central coordinator of neuroinflammatory activation, with both protective and pathologic consequences during neurologic disease states (*21, 22, 45*). Despite this understanding, cell-type specific functions for RIPK3 within the CNS have not been carefully delineated. We have shown previously that neuronal RIPK3 is required for cell-intrinsic expression of antiviral effector genes, as well as for the expression of chemokine molecules that recruit infiltrating leukocytes to the CNS during flavivirus infection, and that both of these mechanisms are required for host protection (*29–31*). In contrast, we recently reported that RIPK3 signaling in astrocytes is deleterious in models of Parkinsonian neurodegeneration, where it promotes pathogenic neuroinflammation and neurotoxicity (*39, 40*). Here, we demonstrate a more nuanced role for astrocytic RIPK3 in regulating neuroinflammation during viral encephalitis. While RIPK3 in astrocytes promoted a robust inflammatory transcriptomic signature, it also coordinated expression of immunomodulatory genes that limit the extent of leukocyte infiltration and immunopathology. These findings underscore the potential for specialized, cell-type specific functions for this pathway in maintaining a balanced and targeted immune response within the CNS.

Our findings also highlight a critical role for astrocytes in the control of protease activity in the CNS, with important implications for BBB integrity and leukocyte recruitment during neuroinflammation. Endogenous proteases, including gelatinases such as MMPs, play significant roles in the pathogenesis of flavivirus encephalitis. MMP-9, for example, is upregulated during flavivirus encephalitis and contributes to BBB breakdown by degrading TJ proteins and extracellular matrix components like collagen (*46–48*). This proteolytic activity facilitates the infiltration of immune cells into the CNS, exacerbating neuroinflammation and tissue damage (*49, 50*). Similarly, granzyme B is a serine protease released by cytotoxic T cells and natural killer cells and is highly upregulated in brain-infiltrating lymphocytes during flavivirus encephalitis (*13, 51–53*). In addition to its role in degrading BBB components, granzyme B is a significant source of immunopathology due its intrinsic neurotoxic activity (*54–57*). These activities highlight the need for tight regulation of inflammatory protease activity to balance effective antiviral immune responses with the preservation of CNS homeostasis.

In the current study, we show that protease regulation through serpins, and particularly SerpinA3N, is essential for maintaining acceptable levels of gelatinase activity and preserving BBB tight junctions during flavivirus encephalitis. These findings are consistent with previous reports showing that SerpinA3N effectively protected mice from excessive neuroinflammation and promoted functional recovery in models of ischemic stroke, traumatic brain injury, and multiple sclerosis (*58–62*). However, the role of SerpinA3N in the CNS appears to be quite complex, with a range of studies reporting both beneficial (*58–63*) and detrimental (*64–67*) effects in different disease contexts. This complexity can probably be partially attributed to the diverse interaction partners of SerpinA3N, which include a broad array of serine proteases (*44*). Identifying the specific proteases whose inhibition is key to the disease modifying effects of serpin activity has been a challenge in the literature thus far and is a key future direction for our work. Several other variables, including extracellular vs. intracellular sites of action, may also underlie the differential effects of serpins in distinct disease contexts (*41, 68*). The pleotropic nature of the serine protease targets of SerpinA3N may also underly some of these apparently contradictory findings; for example, target proteases such as ADAM10/17, MMP-9, and the tPA/plasmin system have major roles in promoting inflammatory responses, but also fulfill critical homeostatic functions in the CNS (*69–73*). The impact of SerpinA3N and related protease inhibitors is thus likely influenced by the context of their expression and the relative abundance, localization, and functional status of the proteases they target across diverse disease states. Clarifying these interactions will be vital for distinguishing between the beneficial and harmful effects of proteases and their inhibitors in CNS health and disease.

Our study also adds to a growing awareness of the pathogenic role of CD8^+^ T cells in the brain during viral encephalitis. Past work suggests that some amount of cytotoxic lymphocyte response in the brain is required for viral clearance and host survival during flavivirus encephalitis (*74–76*). However, interventions that suppress CD8^+^ T cell infiltration and activity in the brain have been shown, paradoxically, to improve host survival and ameliorate neuropathogenesis, without sacrificing virologic control (*18, 19, 35*). T cell responses in the CNS are under complex regulation, and the balance of their protective vs. pathogenic activities is also likely highly context dependent. For example, past work has shown that release of CD8^+^ T cells from perivascular spaces via CXCR4 antagonism facilitated their access to parenchymal antigen, which improved viral clearance and actually diminished immunopathology (*77*). Similarly, the activation state of antigen presenting cells within the CNS influences the local reactivation of antigen-specific lymphocytes, optimizing their effector function to promote virologic control and minimize off target effects on sensitive bystander cells (*52*). We speculate that the activity of SerpinA3N may influence lymphocyte recruitment in ways that preserve CD8^+^ T cell-mediated protection while suppressing the recruitment or aberrant activation of neuropathogenic lymphocytes. This would represent an intriguing mechanism of cross-talk between neurons and astrocytes during flavivirus encephalitis, with neuronal RIPK3 signaling largely promoting CD8^+^ T cell recruitment while astrocytic RIPK3 limits and/or optimizes this response to promote neuroprotection over neuropathogenesis.

## Methods

### Mouse lines

All mice in this study were bred and housed under specific-pathogen free conditions in Nelson Biological Laboratories at Rutgers University. All lines were backcrossed to be congenic to the C57BL/6J background (Jackson Laboratories, #000664). Adult (>8 week old) animals of both sexes were used in all studies, following protocols approved by the Rutgers University Institutional Animal Care and Use Committee (IACUC). *Ripk3*^−/−^ and *Ripk3*^fl/fl^ mouse lines were generously provided by Genentech, Inc (San Francisco, CA, USA). *Mlkl*^−/−^ (*78*) and *Ripk3*-2xFV^fl/fl^ (*30*) lines were provided by Andrew Oberst (University of Washington, Seattle, WA, USA). *Aldh1l1*-Cre/ERT2 mice were obtained from Jackson Laboratories (#031008) and all animals expressing this transgene were treated for five days with 60 mg/kg tamoxifen (Sigma-Aldrich, #T5648) in sunflower oil (Sigma-Aldrich, #S5007) (i.p.) at least one week prior to further experimentation.

### Viruses and virologic assays

ZIKV-MR766 was provided by the World Reference Center for Emerging Viruses and Arboviruses (WRCEVA). ZIKV-Fortaleza was provided by Andrew Oberst (University of Washington, Seattle, WA, USA). WNV strain WN02-Bird 114 (*79*) was generously provided by Dr. Bobby Brooke Hererra (Rutgers Robert Wood Johnson Medical School, New Brunswick, NJ, USA). Viral stocks were generated by infecting Vero cells (ATCC, #CCL-81) at an MOI of 0.01 and harvesting supernatants at 72hpi. Viral titers of stocks were determined via plaque assay on Vero cells. Cells were maintained in DMEM (Corning #10–013-CV) supplemented with 10% Heat Inactivated FBS (Gemini Biosciences #100–106), 1% Penicillin–Streptomycin-Glutamine (Gemini Biosciences #400–110), 1% Amphotericin B (Gemini Biosciences #400–104), 1% Non-Essential Amino Acids (Cytiva, #SH30238.01), and 1% HEPES (Cytiva, #SH30237.01). Plaque assay media was composed of 1X EMEM (Lonza # 12–684F) supplemented with 2% Heat Inactivated FBS (Gemini Biosciences #100–106), 1% Penicillin–Streptomycin-Glutamine (Gemini Biosciences, #400–110), 1% Amphotericin B (Gemini Biosciences #400–104), 1% Non-Essential Amino Acids (Cytiva, #SH30238.01), and 1% HEPES (Cytiva, SH30237.01), 0.75% Sodium Bicarbonate (VWR, #BDH9280) and 0.5% Methyl Cellulose (VWR, #K390). Plaque assays were developed at 4 days post infection by removal of overlay medium and staining/fixation using 10% neutral buffered formalin (VWR, #89370) and 0.25% crystal violet (VWR, #0528). Plaque assays were performed by adding 100uL of serially diluted sample for 1 hour at 37°C to 12-well plates containing 200,000 Vero cells per well. Plates were further incubated with plaque assay media at 37°C and 5% CO_2_ for 5 days. Medium was removed from the wells and replaced with fixative containing crystal violet for approximately 20–30 minutes. Plates were washed repeatedly in H_2_O and allowed to dry before counting visible plaques.

### Mouse infections and tissue harvesting

Isoflurane anesthesia was used for all procedures. Unless otherwise noted, mice were inoculated subcutaneously (50uL) with 3×104 PFU WNV-WN02-Bird114 or intracranially (10uL) with 50 PFU ZIKV-MR766/ZIKV-Fortalea. At appropriate times post infection, mice underwent cardiac perfusions with 30 mL cold sterile 1X phosphate-buffered saline (PBS). Extracted tissues were weighed and homogenized using 1.0 mm diameter zirconia/silica beads (Biospec Products, #11079110z) in sterile PBS for plaque assay or TRI Reagent (Zymo, #R2050–1) for gene expression analysis. Homogenization was performed in an Omni Beadrupter Elite for 2 sequential cycles of 20 s at a speed of 4 m/s.

### Primary cell culture

Primary cortical astrocytes were generated from P1-P3 mouse pups, as previously described (*39*). Tissues were dissociated using the Neural Dissociation Kit (T) following manufacturer’s instructions (Miltenyi, #130–093-231). Astrocytes were expanded in AM-a medium (ScienCell, #1831) supplemented with 10% FBS in fibronectin-coated cell culture flasks and seeded into plates coated with 20 μg/mL Poly-L-Lysine (Sigma-Aldrich, #9155) before experiments.

### Quantitative real-time PCR

Total RNA from harvested tissues was extracted using Zymo Direct-zol RNA Miniprep kit, as per manufacturer instructions (Zymo, #R2051). Total RNA extraction from cultured cells, cDNA synthesis, and subsequent qRT-PCR were performed as previously described (*39, 80*). Cycle threshold (CT) values for analyzed genes were normalized to CT values of the housekeeping gene 18 S (CT_Target_ - CT_18S_ = ΔCT). Data from primary cell culture experiments were further normalized to baseline control values (ΔCT_experimental_ - ΔCT_control_ = ΔΔCT (DDCT)). Primer sequences for qRT-PCR analysis can be found in Table S2.

### Bulk RNA sequencing

*Ripk3*-2XFV^fl/fl^ *Aldh1l1* Cre^+^ mice and Cre^−^ littermate controls were treated intraperitoneally with B/B Homodimerizer (Takara, #AP20187) for 24 hours prior to perfusion with PBS and harvesting of whole brain. Brains were digested and myelin removed using the Adult Brain Dissociation Kit (Miltenyi, #130-107-677). Astrocytes were isolated using the Anti-ACSA-2 MicroBead Kit (Miltenyi, #130-097-678) and RNA extracted using RNeasy Micro Kit (Qiagen, # 74004). RNA library preparation, sequencing, and preliminary analysis was performed as described (*81*) by Azenta Life Sciences. The GEO accession number for this dataset will be available upon publication.

### Flow Cytometry

Mouse brains were dissected from freshly perfused mice and placed into tubes containing 1X PBS. Brain tissues were incubated with 10mL buffer containing 0.05% Collagenase Type I (Sigma-Aldrich, #C0130), 10ug/mL DNase I (Sigma-Aldrich, #D4527) and 10mM HEPES (Cytiva, #SH30237.01) in 1X Hanks’ Balanced Salt Solution (VWR, #02–1231-0500) for one hour at room temperature under constant rotation. Brain tissues were transferred to a 70um strainer on 50mL conical tubes and mashed through the strainer using the plunger of 3–5mL syringes. Tissue was separated in 8 mL 37% Isotonic Percoll (Percoll: Cytiva, #17–0891-02; RPMI 1640: Corning, #10–040-CV, supplemented with 5% FBS) by centrifugation at 1200xg for 30 minutes with a slow brake. The myelin layer and supernatant were discarded. Leukocytes were incubated in 1X RBC Lysis Buffer (Tonbo Biosciences, #TNB-4300-L100) for 10 minutes at room temperature. Cells were centrifuged and resuspended in FACS buffer composed of 1X PBS, 0.2% sodium azide and 5% FBS. Samples were transferred into a U-bottomed 96-well plate. Leukocytes were blocked with 2% normal mouse serum and 1% FcX Block (BioLegend, #101320) in FACS buffer for 30 minutes at 4°C prior to being stained with Zombie NIR (Biolegend, #423105) for 15 minutes at room temperature. Cells were then stained with fluorescently conjugated antibodies to CD3e (Biolegend, clone 17A2), CD44 (Biolegend, clone IM7), CD8a (Biolegend, clone 53–6.7), CD4 (Biolegend, clone RM4–5), CD45.2 (Biolegend, clone 104), MHC-II (Biolegend, clone M5/114.15.2), NK1.1 (Biolegend, clone PK136), CD11c (Biolegend, clone N418), F4/80 (Biolegend, clone BM8), CD11b (Biolegend, clone M1/70), Ly6G (Biolegend, clone 1A8), Ly6C (Biolegend, clone HK1.4), and CD80 (Biolegend, clone 16–10A1). Leukocytes were stained for 30 minutes at 4C prior to washing in FACS buffer and fixation with 1% PFA in PBS (ThermoFisher, #J19943-K2). Data collection and analysis were performed using a Cytek Northern Lights Cytometer (Cytek) and FlowJo software (Treestar). Data were normalized using a standard bead concentration counted by the cytometer with each sample (ThermoFisher, #C36950). Spleens were crushed between two slides, filtered through a 70um cell strainer, and washed with FACS buffer. Isolated splenocytes were incubated with 1X RBC Lysis Buffer as done for leukocytes isolated from the brain prior to blocking and staining.

### CD8 Depletion

Mice were injected intraperitoneally with 450μg of anti-CD8 antibody (ThermoFisher, clone H35-17.2). Blood samples were collected retro-orbitally two days later and circulating leukocytes assessed by flow cytometry to confirm depletion of CD8^+^ cells.

### SerpinA3N treatment

Mice were injected ICV with 40ng recombinant mouse SerpinA3N (R&D Systems, # 4709-PI-010) diluted in HBSS. Primary astrocyte cultures were treated with 40ng of recombinant mouse SerpinA3N one hour before infection for virologic assays.

### Gelatinase/collagenase activity assay

EnzCheck Gelatinase/Collagenase Assay Kit (ThermoFisher, E12055) was used according to manufacturer’s protocol. Mice were perfused at four days post infection using cold sterile 1XPBS, followed by harvesting of the brain. A sagittal cut divided the brain in two halves, one of which was homogenized in 1X reaction buffer. Brian homogenate was centrifuged at 15,000 x g for 10 minutes at 4C and clarified supernatants were used for the assay. Samples were incubated with fluorogenic gelatin substrate in a black-walled 96-well plate for two hours at room temperature. Samples were read on a microplate reader at 485/535nm.

### SerpinA3N ELISA

SerpinA3N protein concentrations in brain tissue homogenates were measured using a mouse SerpinA3N ELISA kit (RayBiotech, ELM-SERPINA3N-1) according to manufacturer’s instructions.

### Immunofluorescent imaging

For imaging of tight junctions, mice were perfused with 30mL 1X PBS followed by cold methanol. Brains were placed in methanol overnight and rehydrated in 1X PBS. For other stains, mice were perfused with 30mL cold 1XPBS followed by freshly prepared 4% paraformaldehyde in 1X PBS. Brains were stored at 4C overnight, followed by replacement of PFA with 1X PBS until sectioning was performed. In all cases, brains were embedded in agar and sectioned using a compresstome (Presisionary, VF-510-0Z). Sections were blocked with 10% goat or donkey serum with 0.4% Triton X-100 followed by incubation with primary antibodies for 48 hours at 4C. Antibodies used were NeuN (1:500, Synaptic systems, 266-004), Pan-flavivirus E protein antibody (1:100, VWR Enzo, 76285-044), GFAP (1:250, Invitrogen, 13-0300), Claudin-5 (1:200, Invitrogen, 34-1600), and Zo-1 (1:200, Millipore, MABT11). Sections were washed prior to incubation for 1 hour at room temperature with secondary antibodies. Sections were stained with DAPI prior to mounting with ProLong Diamond AntiFade (ThermoFisher, P36970).

### Statistical analysis

Statistical analyses were performed using GraphPad Prism 9. Survival experiments were compared via log-rank test. Most other experiments were compared with appropriate parametric tests, including the Student’s t-test (two-tailed) or two-way analysis of variance (ANOVA) with Tukey’s post hoc test to identify significant differences between groups. A p-value of less than 0.05 was deemed to indicate statistical significance. Unless specified otherwise, all data points represent biological replicates consisting of distinct mice or independent cultures derived from distinct mice.

## Acknowledgements

This work was supported by R01 NS120895 (to BPD) and R00 HD103911 (to NMO). IE was supported by an HHMI Gilliam Fellowship.

## Author Contributions

Conceptualization: ML, BPD; Investigation: ML, IE, EM, KN, CA, NMO, BPD; Analysis: ML, IE, EM, EMD, TWC, NMO, BPD; Resources: NMO, BPD; Writing – Original Draft: ML, EM, BPD; Writing – Review and Editing: ML, CA, NMO, BPD; Supervision: CA, NMO, BPD; Funding Acquisition: NMO, BPD.

## Supplementary Materials

**Figure S1.**
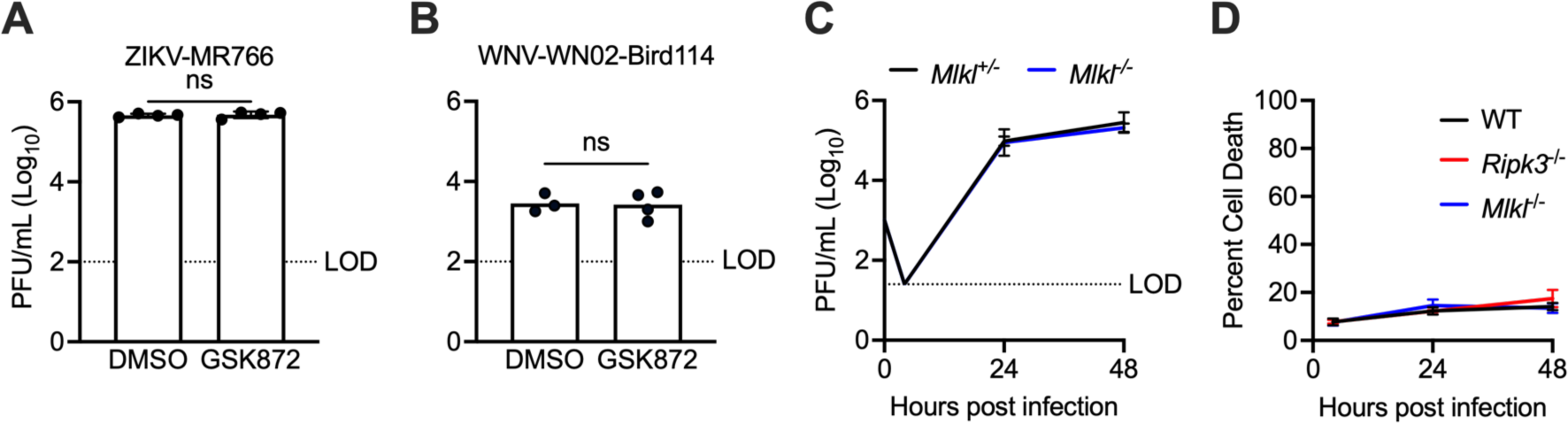
(**A-B**) Viral titer measurements via plaque assay of primary mouse astrocyte supernatants 24 hours following 0.1 MOI ZIKV-MR766 infection (A) or 0.01 MOI WNV-WN02-Bird114 infection (B). Cultures were pretreated with GSK872 for DMSO vehicle two hours prior to infection. (**C**) Multistep growth curve analysis of ZIKV-MR766 replication in primary cortical astrocytes derived from mice of indicated genotypes. Viral titers were assessed via plaque assay. N=4 replicates/group. (**D**) Cell viability measured via Cell Titer Glo assay in primary cortical astrocytes of indicated genotypes following infection with ZIKV-MR766. Error bars represent SEM.

**Table S1.** Gene ontology enrichment analysis of terms related to immune cell trafficking and inflammatory signaling in ACSA-2^+^ astrocytes following chemogenetic RIPK3 activation as described in Figure 2C.

| Gene Ontology Term | Number of Genes | Fold Enrichment | FD Rate (adj. P-value) |
| --- | --- | --- | --- |
| Neutrophil chemotaxis | 33 | 2.91 | 8.01E-05 |
| T cell mediated immunity | 22 | 2.62 | 7.23E-03 |
| Transport across blood-brain barrier | 12 | 4.07 | 8.36E-03 |
| Leukotriene metabolic process | 12 | 3.54 | 1.85E-02 |
| Vasodilation | 23 | 2.36 | 2.14E-02 |
| Positive regulation of acute inflammatory response | 20 | 2.38 | 2.94E-02 |
| Positive regulation of cytokine response | 22 | 2.33 | 2.94E-02 |
| Response to type I interferon | 16 | 2.71 | 3.14E-02 |
| Leukocyte Aggregation | 9 | 4.07 | 3.38E-02 |
| Positive regulation of leukocyte differentiation | 51 | 1.64 | 3.52E-02 |
| Process and present peptide antigens via MHC class I | 20 | 2.3 | 3.61E-02 |
| Positive regulation of fever generation | 7 | 5.27 | 3.71E-02 |
| Positive regulation of myeloid mediated immunity | 19 | 2.34 | 4.06E-02 |
| Amyloid-beta clearance | 11 | 3.24 | 4.12E-02 |
| Maintenance of blood-brain barrier | 10 | 3.39 | 4.80E-02 |
| Positive chemotaxis | 10 | 3.39 | 4.81E-02 |

**Figure S2.**
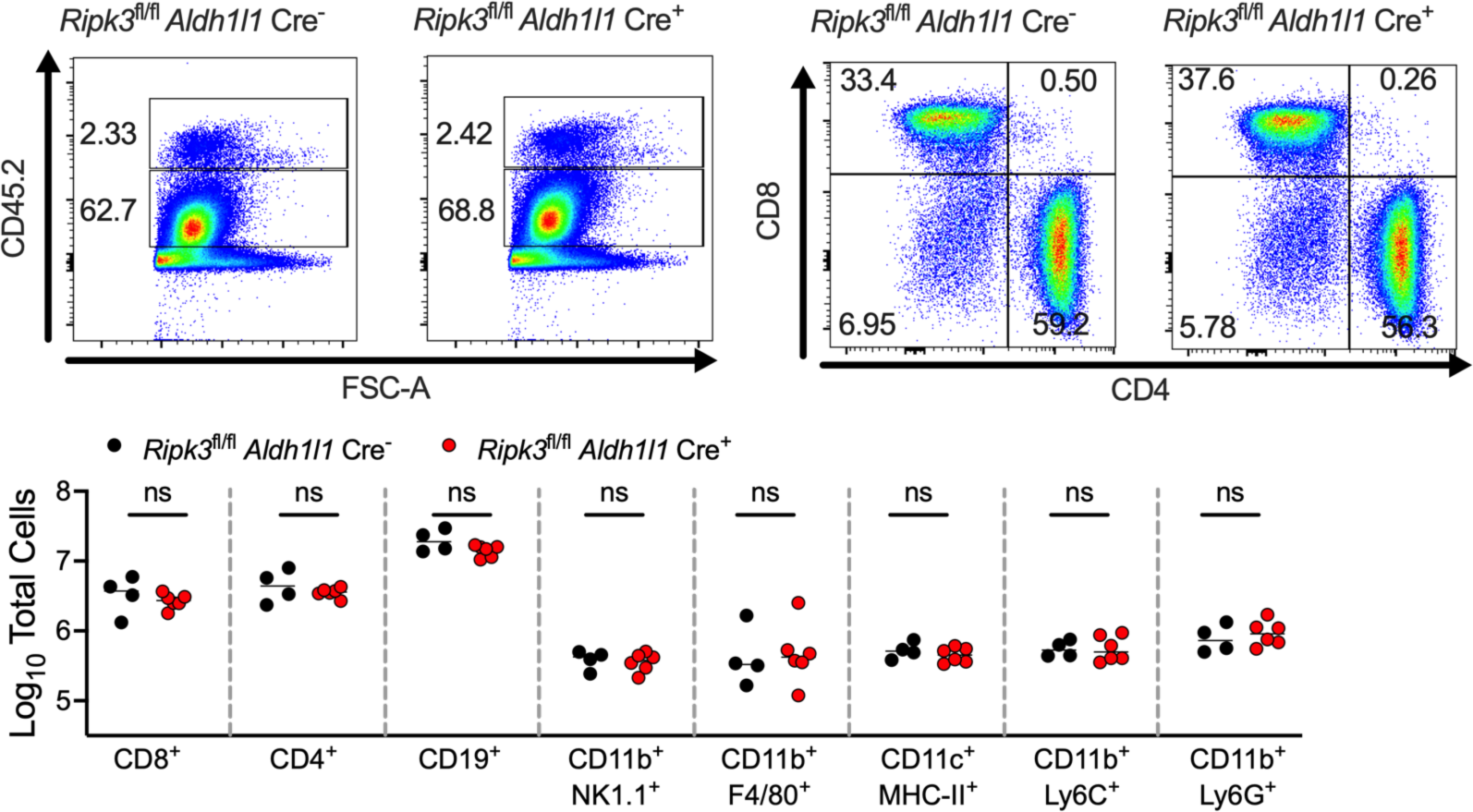
(**A**) Representative flow cytometry plots depicting resident (CD45.2^int^) or infiltrating (CD45.2^hi^) CNS leukocytes derived from mice of indicated genotypes four days following mock intracranial infection (injection of HBSS). (**B**) Representative flow cytometry plots depicting splenic CD4^+^ T cells and CD8^+^ T cells derived from mice of indicated genotypes four days following ZIKV-MR766 infection. (**C**) Total numbers of indicated leukocyte populations in spleen tissues derived from mice of indicated genotypes four days following ZIKV-MR766 infection.

**Figure S3.**
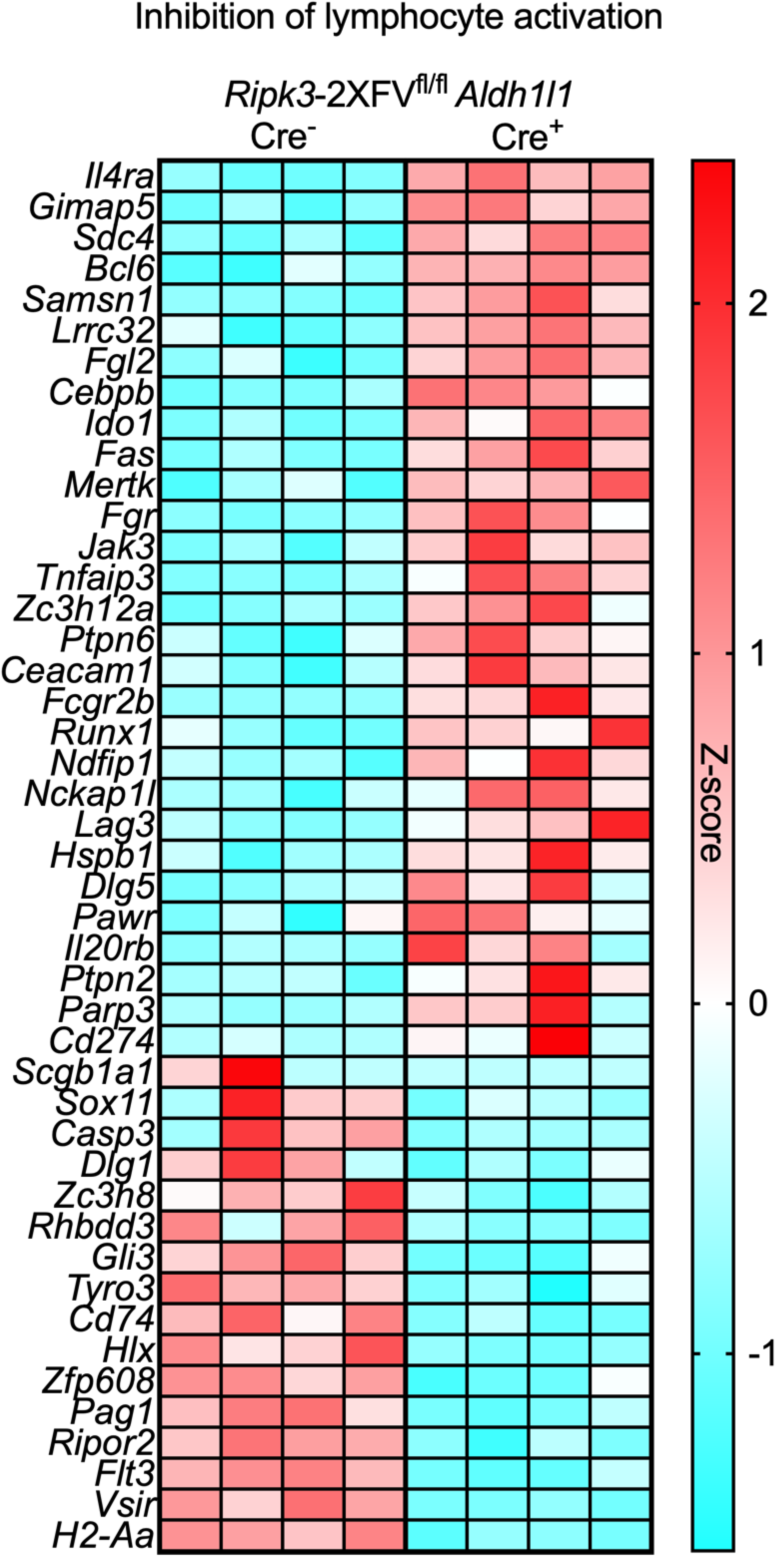
Heatmap depicting significant differentially expressed genes within indicated gene ontology term. Data derived from RNAseq analysis comparing ACSA-2^+^ astrocytes sorted from *Ripk3*-2xFV^fl/fl^ *Aldh1l1* Cre^+^ mice compared to astrocytes sorted from Cre^−^ littermate controls 24 hours following B/B treatment.

**Figure S4.**
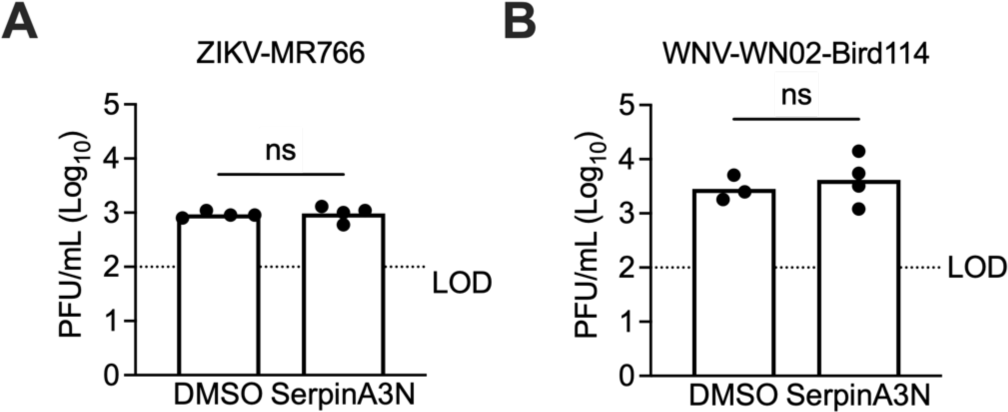
(**A-B**) Viral titer measurements via plaque assay of primary mouse astrocyte supernatants 24 hours following 0.01 MOI ZIKV-MR766 infection (A) or 0.01 MOI WNV-WN02-Bird114 infection (B). Cultures were pretreated with SerpinA3N or DMSO vehicle two hours prior to infection. LOD=limit of detection.

**Figure S5.**
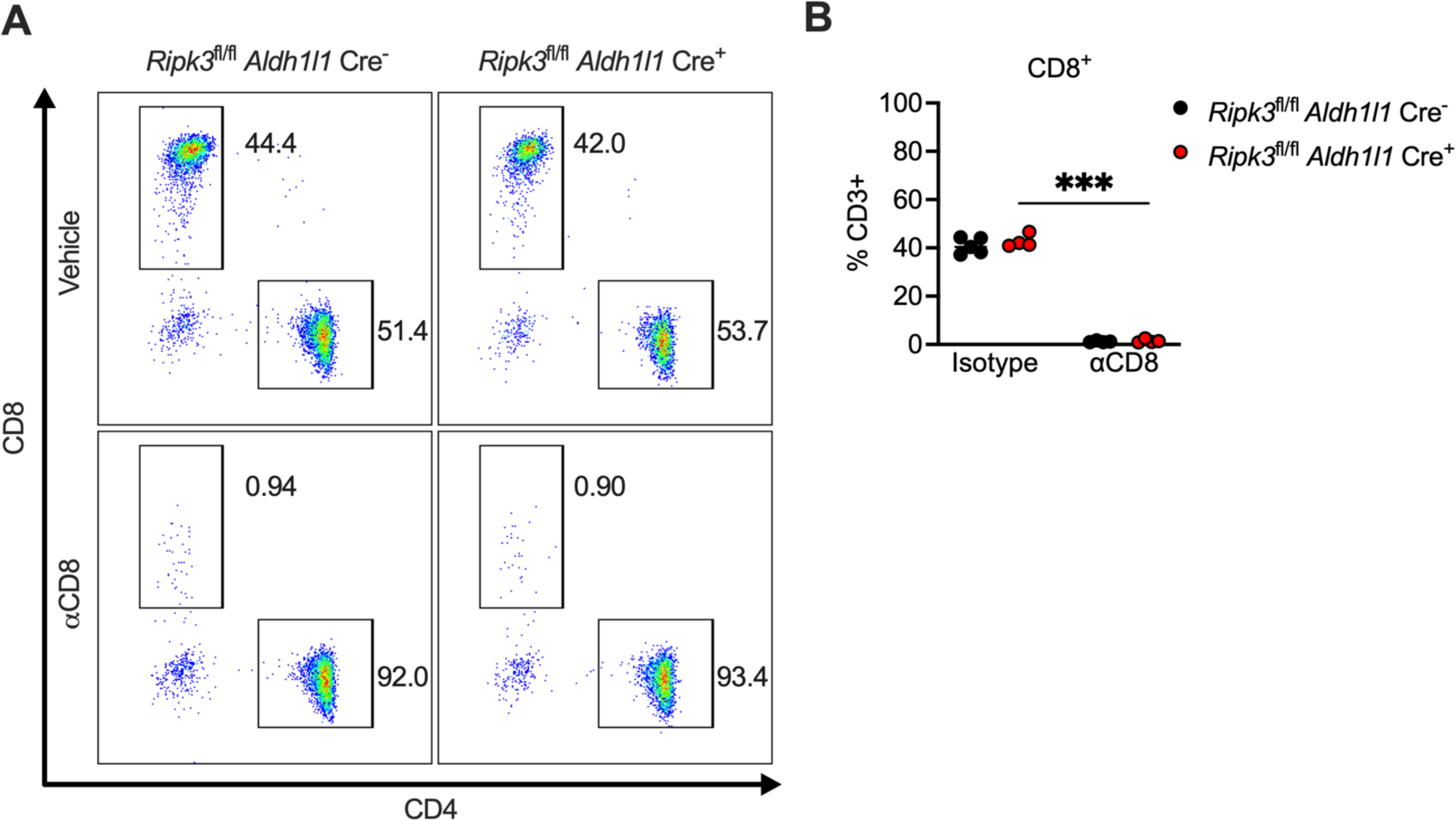
(**A**) Representative flow cytometry plots depicting CD4^+^ T cells and CD8^+^ T cells in blood samples from mice of indicated genotypes. Mice were infected intracranially with ZIKV-MR766, followed by intraperitoneal administration of an anti-CD8 neutralizing antibody (αCD8) or isotype control two days post infection. Flow cytometry was performed at four days post infection. (**B**) Total numbers of CD8^+^ T cells in blood samples of mice described in (A). ***p < 0.001.

**Table S2:**
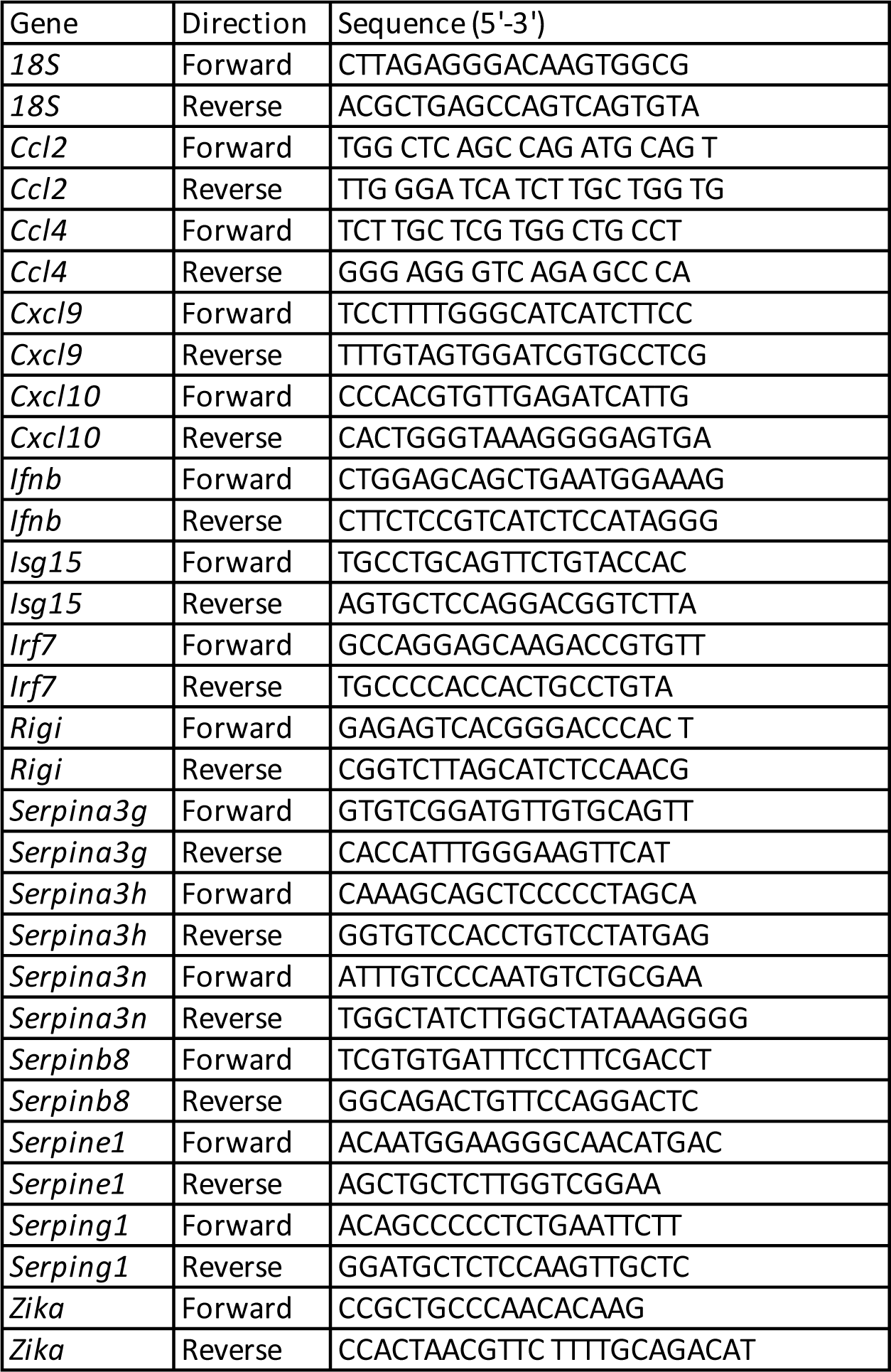
Primer sequences for qRT-PCR.

